# Screening for neurotoxic potential of 15 flame retardants using freshwater planarians

**DOI:** 10.1101/538280

**Authors:** Siqi Zhang, Danielle Ireland, Nisha S. Sipes, Mamta Behl, Eva-Maria S. Collins

## Abstract

Asexual freshwater planarians are an attractive invertebrate model for high-throughput neurotoxicity screening, because they possess multiple quantifiable behaviors to assess distinct neuronal functions. Planarians uniquely allow direct comparisons between developing and adult animals to distinguish developmentally selective effects from general neurotoxicity. In this study, we used our automated planarian screening platform to compare the neurotoxicity of 15 flame retardants (FRs), consisting of representative phased-out brominated (BFRs) and replacement organophosphorus FRs (OPFRs). OPFRs have emerged as a proposed safer alternative to BFRs; however, limited information is available on their health effects. We found 11 of the 15 FRs (3/6 BFRs, 7/8 OPFRs, and Firemaster 550) caused adverse effects in both adult and developing planarians with similar nominal lowest-effect-levels for BFRs and OPFRs. This suggests that replacement OPFRs are comparably neurotoxic to the phased-out compounds. BFRs were primarily systemically toxic, whereas OPFRs, except Tris(2-chloroethyl) phosphate, shared a behavioral phenotype in response to noxious heat at sublethal concentrations, indicating specific neurotoxic effects. By directly comparing effects on adult and developing planarians, we further found that one BFR (3,3’,5,5’-Tetrabromobisphenol A) caused a developmental selective defect. Together, these results demonstrate that our planarian screening platform yields high content data resulting from assaying various behavioral and morphological endpoints, allowing us to distinguish selective neurotoxic effects and effects specific to the developing nervous system. Ten of these 11 bioactive FRs were previously found to be bioactive in other models, including cell culture and alternative animal models (nematodes and zebrafish). This level of concordance across different platforms emphasizes the urgent need for further evaluation of OPFRs in mammalian systems.

**Abbreviations:**

DNT: developmental neurotoxicity
FR: flame retardant
BFR: brominated flame retardant
PBDE: Polybrominated Diphenyl Ether
OPFR: organophosphorus flame retardant
NTP: National Toxicology Program
DMSO: dimethyl sulfoxide
HTS: high-throughput screening
LEL: lowest effect level
HTTK: high-throughput toxicokinetics
POD: point of departure

## 1. Introduction

Traditional hazard assessment using mammalian models cannot keep up with the speed of chemical development. The Toxic Substances Control Act Inventory lists approximately 68,000 chemicals currently in use in the United States (https://www.epa.gov/tsca-inventory), but <2% have been evaluated for developmental neurotoxicity (DNT) (Smirnova et al., 2014). *In vitro* testing combined with bioinformatic analyses have emerged as a powerful tool to fill this need for more rapid chemical testing (Fritsche et al., 2018), but this approach faces limitations for predicting adverse effects on neuronal function (Smirnova et al., 2014; Tohyama, 2016). Non-mammalian animal models, amenable to high-throughput, cost-effective screening, are a promising alternative solution for efficient DNT screening since they allow for automated behavioral assays to quantitatively evaluate different neuronal functions (Bal-Price et al., 2012).

Flame retardants (FRs), widely added to commercial products for fire safety (Costa and Giordano, 2007; Salimi et al., 2017), are a class of chemicals which only have limited mammalian data for safety assessment. Prior to 2005, brominated FRs (BFRs) such as polybrominated diphenyl ether (PBDE) mixtures were the primary FRs used in the United States (Hale et al., 2003). Since then, many were voluntarily phased-out due to growing evidence of their associations with impaired neurodevelopment and decreased fertility, exacerbated by their persistence in the environment and ability to bioaccumulate (Darnerud, 2003; Herbstman et al., 2010; Stapleton et al., 2011; Talsness, 2008). Organophosphorus FRs (OPFRs) have gradually become replacements for PBDEs over the last decade (Stapleton et al., 2012; Stapleton et al., 2014). Concerns exist that OPFRs, similar to PBDEs, may persist in the environment and bioaccumulate, but little is known about their potential toxicity (Hendriks and Westerink, 2015; Meeker and Stapleton, 2010; Stapleton et al., 2009; Van Der Veen and De Boer, 2012). Particularly concerning is the potential DNT of OPFRs, as they share structural similarities to organophosphorus pesticides, which adversely affect neurodevelopment and cause neurobehavioral impairments (González-Alzaga et al., 2014; Muñoz-Quezada et al., 2013; Ricceri et al., 2006; Slotkin et al., 2006). Previous work using a screening battery of *in vitro* and alternative animal models found that several OPFRs had comparable toxicity and potency to BFRs, including phased-out BDE-47 (Behl et al., 2015; Behl et al., 2016; Jarema et al., 2015), indicating that OPFRs may not be the safer alternative to BFRs that they have been marketed to be.

Because the freshwater planarian *Dugesia japonica* is a good model to study organophosphorus pesticide neurotoxicity (Hagstrom et al., 2017; Hagstrom et al., 2018a; Zhang et al., 2018) and uniquely allows parallel screening of adult and developing animals to identify developmental-specific effects (Zhang et al., 2018), we herein evaluate the potential neurotoxicity and DNT of 15 FRs using a custom planarian high-throughput screening (HTS) platform. These FRs include 3 phased-out PBDEs (BDE-47, BDE-99, and BDE-153), 3 currently in-use BFRs (TBB, TBBPA, and TBPH), 8 in-use OPFRs (BPDP, EHDP, IDDP, IPP, TCEP, TCPP, TMPP, and TPHP), and one BFR mixture (FM550).

Because *D. japonica* is a fairly new model for neurotoxicity studies, we briefly summarize its key features to familiarize the reader and refer to our recent review paper (Hagstrom et al., 2016) for an in-depth review of planarians as an alternative model for DNT. In asexual *D. japonica* planarians, neurodevelopment is solely achieved through neuroregeneration following asexual reproduction by fission, allowing neurodevelopment to be induced by head amputation. In addition, the similar sizes of adult and regenerating/developing planarians allows for both worm types to be screened in parallel with the same assays (Hagstrom et al., 2015; Hagstrom et al., 2016; Zhang et al., 2018). Although the planarian central nervous system is morphologically simple, consisting of a bilobed cephalic ganglion and two ventral nerve cords, it is molecularly complex and highly compartmentalized. It shares the same major neurotransmitters as the vertebrate brain, including dopamine, serotonin, octopamine, GABA, and acetylcholine (Cebrià, 2007; Cebrià et al., 2002a; Cebrià et al., 2002b; Ross et al., 2017). Furthermore, planarians possess a large repertoire of diverse quantifiable behaviors, such as gliding and swimming, phototaxis, thermotaxis, and scrunching. The molecular mediators of some of these behaviors have been characterized, such as serotonergic neurons in gliding or transient receptor potential channels in thermotaxis (Birkholz and Beane, 2017; Inoue et al., 2014; Nishimura et al., 2010) allowing these behaviors to be used to assess specific neuronal functions, and thus provide insight into the cellular effects of neurotoxicity. In summary, by testing diverse behaviors in both adult and regenerating/developing planarians, this platform uniquely allows differentiation of selective neurotoxic effects and effects specific to the developing nervous system.

The 15 FRs that are studied here comprise a subset of an 87-compound library from the National Toxicology Program (NTP) (Behl et al., 2018; Hagstrom et al., 2018b; Zhang et al., 2018), which we had previously screened. However, in this previous study we had not evaluated these compounds in detail, because we focused on the comparison of compound classes (Zhang et al., 2018) and concordance with other systems (Hagstrom et al., 2018b). Moreover, we re-screened these 15 FRs independently in a second screen and used the two independent screens to evaluate the robustness of our planarian screening platform. Thus, the present study serves a dual purpose: To more closely examine the potential neurotoxicity of FRs and to quantify screen robustness. To contextualize our results, we evaluated concordance to published data from zebrafish, nematode, and human, mouse, and rat cell-culture models (Behl et al., 2015; Behl et al., 2016; Jarema et al., 2015; Noyes et al., 2015). There was high concordance among the different alternative systems showing comparable DNT of OPFRs to BFRs, warranting further investigation into OPFR toxicity.

## 2. Materials and methods

### 2.1. Chemical library

18 chemicals consisting of 6 BFRs (including phased-out BDE-47 and BDE-99), 8 OPFRs, 1 BFR mixture, 2 negative controls, and 1 duplicate (TPHP) were screened in this study. Table 1 lists the 17 unique chemicals with their name, chemical ID, type, supplier, structure, and purity information provided by the NTP (either as determined by the NTP or, if not available, from the chemical supplier’s certificate of analysis). The chemicals were provided by the NTP as ∼20mM stocks in dimethyl sulfoxide (DMSO, Gaylord Chemical Company, Slidell, LA), with the exception of 2,2’,4,4’,5,5’-Hexabromodiphenyl ether (BDE-153), which was provided at 10mM.

**Table 1.**
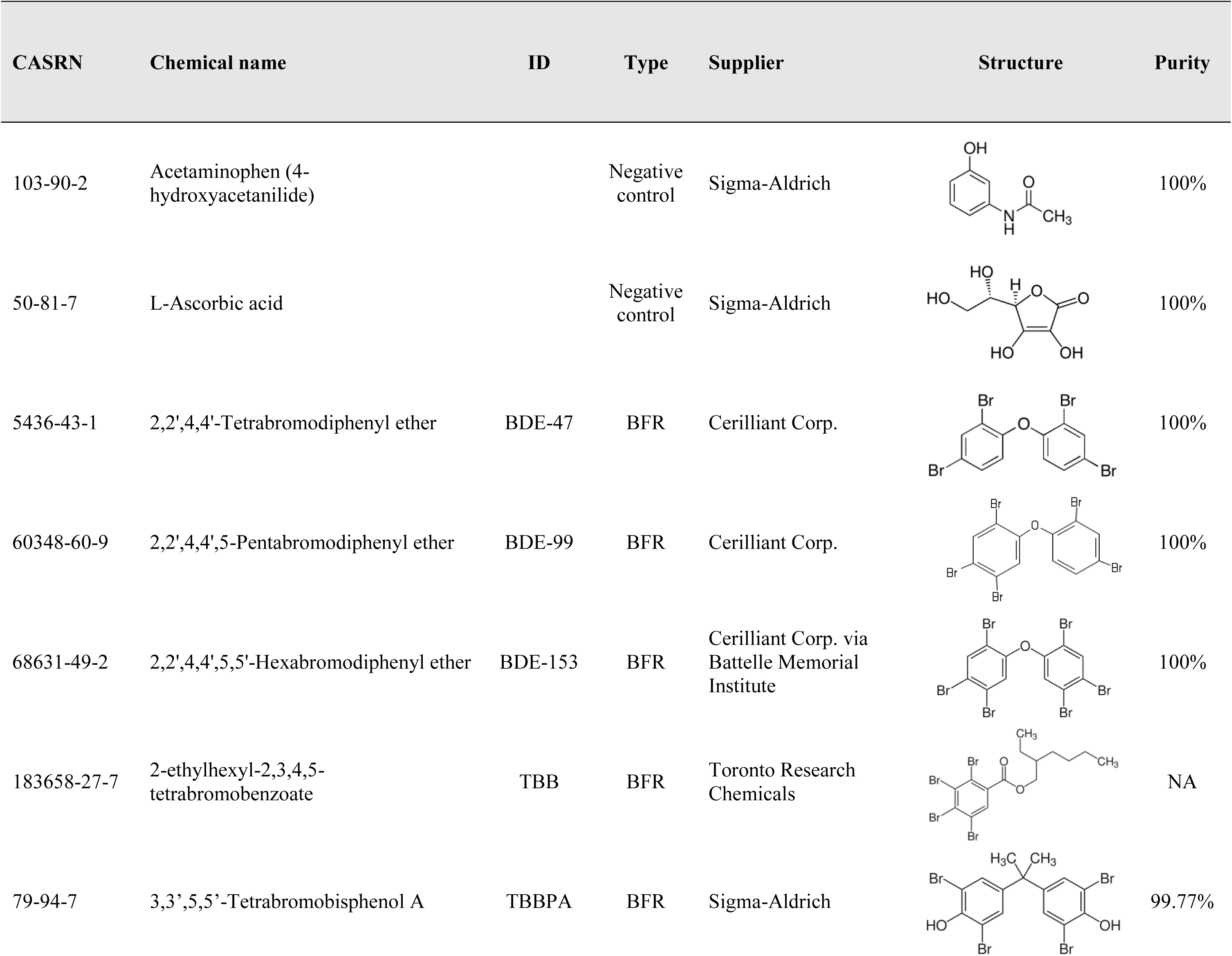
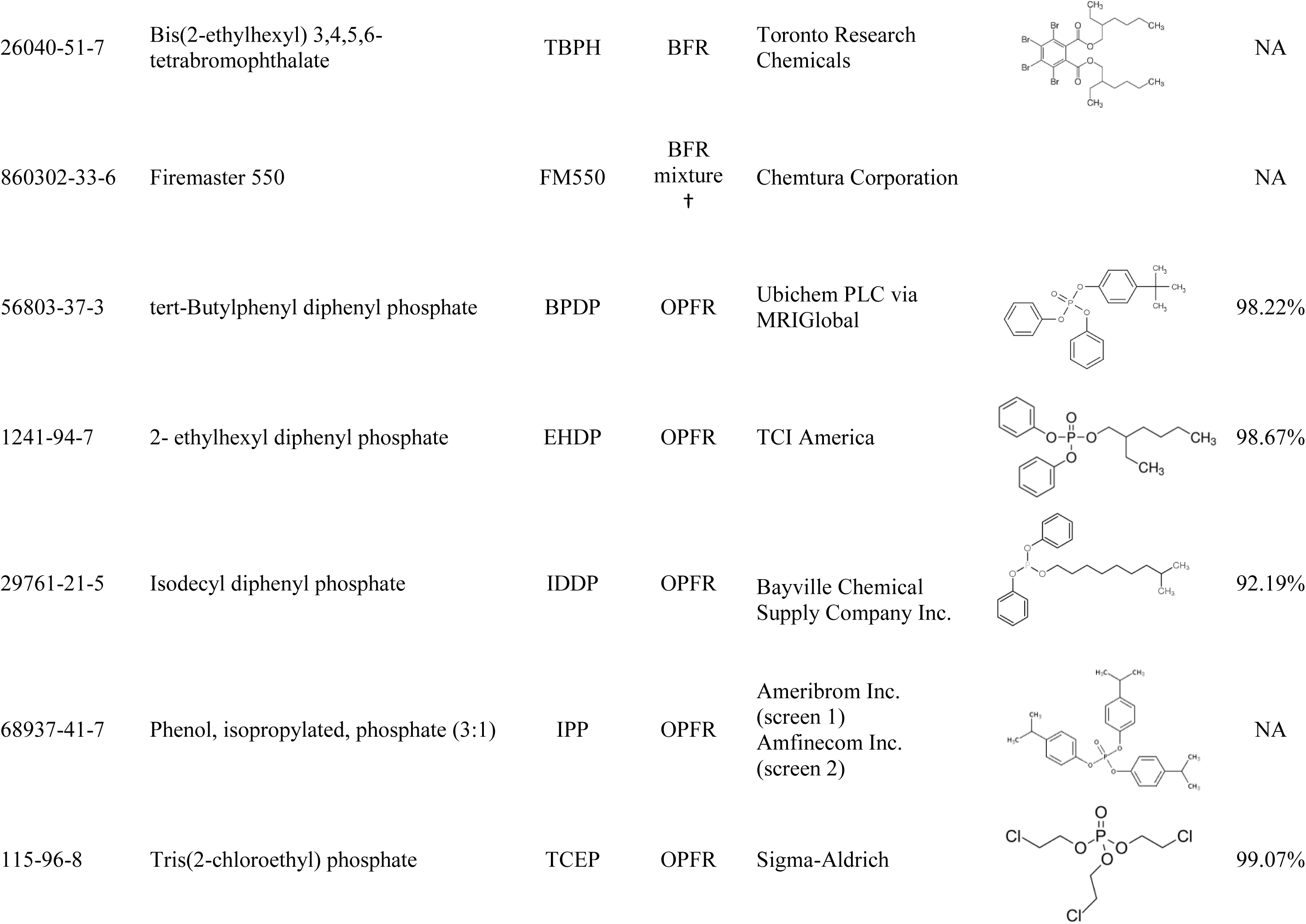
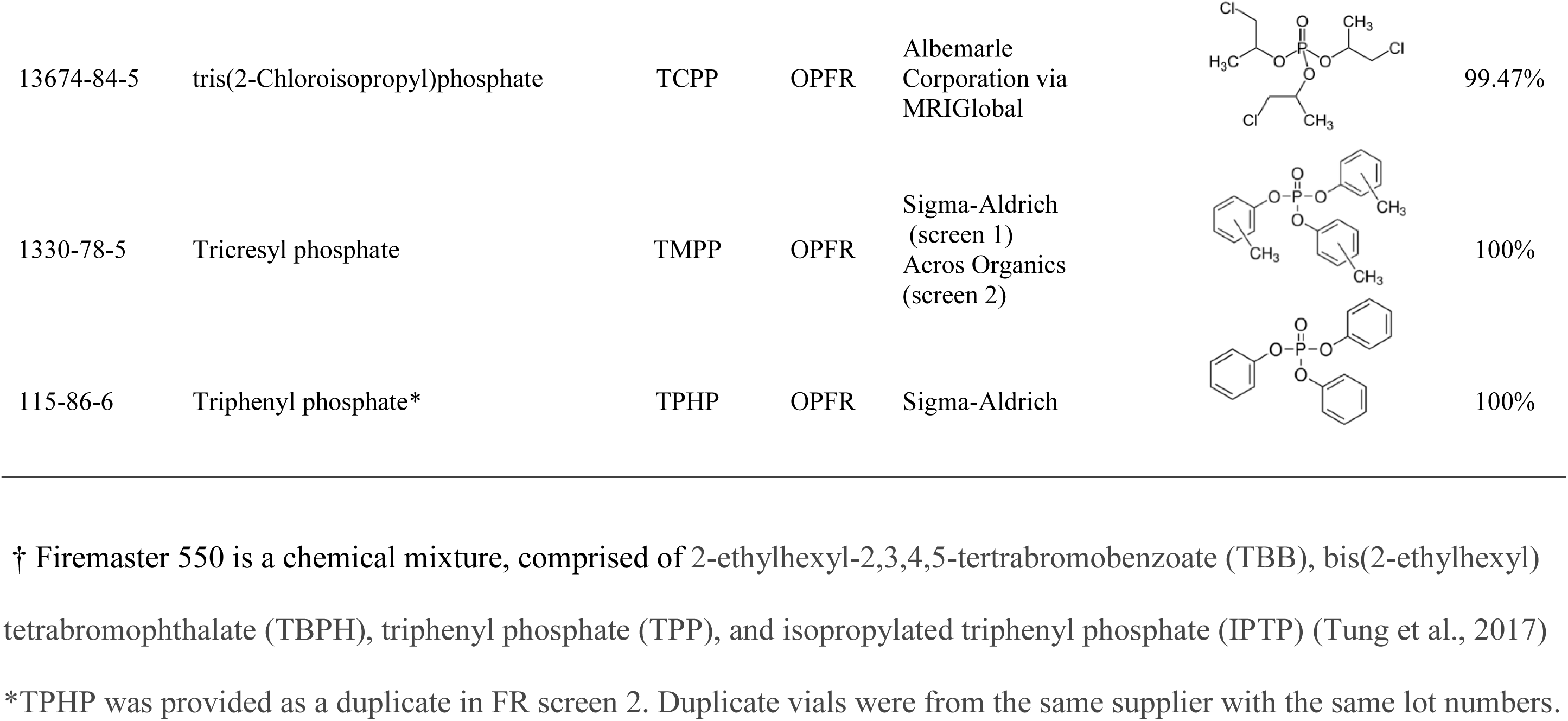
Summary of the screened chemical library with CAS number, chemical name, ID, type (BFR: brominated flame retardant, OPFR: organophosphorus flame retardant), chemical supplier, structure, and purity. When available, the determined purity (by the NTP) is provided, else the manufacturer supplied purity is provided. NA: not available.

| CASRN | Chemical name | ID | Type | Supplier | Structure | Purity |
| --- | --- | --- | --- | --- | --- | --- |
| 103-90-2 | Acetaminophen (4-hydroxyacetanilide) |  | Negative control | Sigma-Aldrich |  | 100% |
| 50-81-7 | L-Ascorbic acid |  | Negative control | Sigma-Aldrich |  | 100% |
| 5436-43-1 | 2,2',4,4'-Tetrabromodiphenyl ether | BDE-47 | BFR | Cerilliant Corp. |  | 100% |
| 60348-60-9 | 2,2',4,4',5-Pentabromodiphenyl ether | BDE-99 | BFR | Cerilliant Corp. |  | 100% |
| 68631-49-2 | 2,2',4,4',5,5'-Hexabromodiphenyl ether | BDE-153 | BFR | Cerilliant Corp. via Battelle Memorial Institute |  | 100% |
| 183658-27-7 | 2-ethylhexyl-2,3,4,5-tetrabromobenzoate | TBB | BFR | Toronto Research Chemicals |  | NA |
| 79-94-7 | 3,3',5,5'-Tetrabromobisphenol A | TBBPA | BFR | Sigma-Aldrich |  | 99.77% |
| 26040-51-7 | Bis(2-ethylhexyl) 3,4,5,6-tetrabromophthalate | TBPH | BFR | Toronto Research Chemicals |  | NA |
| 860302-33-6 | Firemaster 550 | FM550 | BFR mixture<br>† | Chemtura Corporation |  | NA |
| 56803-37-3 | tert-Butylphenyl diphenyl phosphate | BPDP | OPFR | Ubichem PLC via MRIGlobal |  | 98.22% |
| 1241-94-7 | 2-ethylhexyl diphenyl phosphate | EHDP | OPFR | TCI America |  | 98.67% |
| 29761-21-5 | Isodecyl diphenyl phosphate | IDDP | OPFR | Bayville Chemical Supply Company Inc. |  | 92.19% |
| 68937-41-7 | Phenol, isopropylated, phosphate (3:1) | IPP | OPFR | Ameribrom Inc. (screen 1)<br>Amfinecom Inc. (screen 2) |  | NA |
| 115-96-8 | Tris(2-chloroethyl) phosphate | TCEP | OPFR | Sigma-Aldrich |  | 99.07% |
| 13674-84-5 | tris(2-Chloroisopropyl)phosphate | TCPP | OPFR | Albemarle Corporation via MRIGlobal |  | 99.47% |
| 1330-78-5 | Tricresyl phosphate | TMPP | OPFR | Sigma-Aldrich (screen 1)<br>Acros Organics (screen 2) |  | 100% |
| 115-86-6 | Triphenyl phosphate* | TPHP | OPFR | Sigma-Aldrich |  | 100% |

† Firemaster 550 is a chemical mixture, comprised of 2-ethylhexyl-2,3,4,5-tetrabromobenzoate (TBB), bis(2-ethylhexyl) tetrabromophthalate (TBPH), triphenyl phosphate (TPP), and isopropylated triphenyl phosphate (IPTP) (Tung et al., 2017)
\*TPHP was provided as a duplicate in FR screen 2. Duplicate vials were from the same supplier with the same lot numbers.

The tested FRs were a subset of the NTP 87-compound library, which we previously reported on (FR screen 1; (Zhang et al., 2018)). The NTP later provided us with another set of the library (now including 4 duplicate chemicals for a total of 91 chemicals), in which we only screened the subset of chemicals mentioned in this study (FR screen 2). Chemicals in the new library were provided as individual vials, stored at 4ºC and used within 2 months. For all but two chemicals (IPP and TMPP), the same supplier and mostly (14/17) the same chemical lot was used for the chemicals in both FR screen 1 and 2. TPHP was tested in duplicate in FR screen 2, using 2 separate stock vials from the same supplier and lot. The stocks of the negative control L-ascorbic acid were from different lots, but had the same supplier with the same concentration and purities. Five of the chemical stocks differed slightly in concentration between the two libraries, but by at most 0.3mM. Additional information on the NTP library can be found in (Behl et al., 2018).

The chemicals were tested at 0.01, 0.1, 1, 10, and 100μM (0.005, 0.05, 0.5, 5, and 50μM for BDE-153), with a final DMSO concentration of 0.5% for all concentrations. Solvent control populations were exposed to 0.5% DMSO. Chemical dilutions for both libraries were done as described in (Zhang et al., 2018).

### 2.2. FR screen in the planarian system

Asexual *D. japonica*, originally obtained from Shanghai University (Shanghai, China) and cultivated in our lab for > 5 years, were used for all experiments. We screened the library with a custom, fully automated HTS platform using adult (intact) and regenerating, i.e. tail pieces regenerating a new head, planarians in parallel with the same assays, assessing morphological and behavioral endpoints at day 7 and day 12 (Figure 1). Two independent screens, the original NTP 87-compound library screen (“FR screen 1”) and the second, FR-only screen (“FR screen 2”), were performed with 3 replicates (3 independent runs on different days) for each screen. For each replicate, n=8 regenerating and adult planarians were each screened for every chemical concentration. One adult or regenerating animal was loaded into each well of a 48-well plate (Genesee Scientific, San Diego, CA). Static exposure began on day 1, within 3 hours of amputation of the regenerating planarians, by adding the appropriate chemical concentration to the planarians’ aquatic environment. A solvent control population (n=8) exposed to 0.5% DMSO was included in every plate. This DMSO concentration has no effect on planarian morphology or behavior (Hagstrom et al., 2015). The planarians were kept in the dark in the sealed 48-well plates for 12 days, with screening on days 7 and 12 (Figure 1). By day 12, regenerating planarians are considered fully regenerated and behave like adults (Hagstrom et al., 2015; Zhang et al., 2018), thus the two screening days allow us to distinguish regeneration delays from regeneration defects.

**Figure 1.**
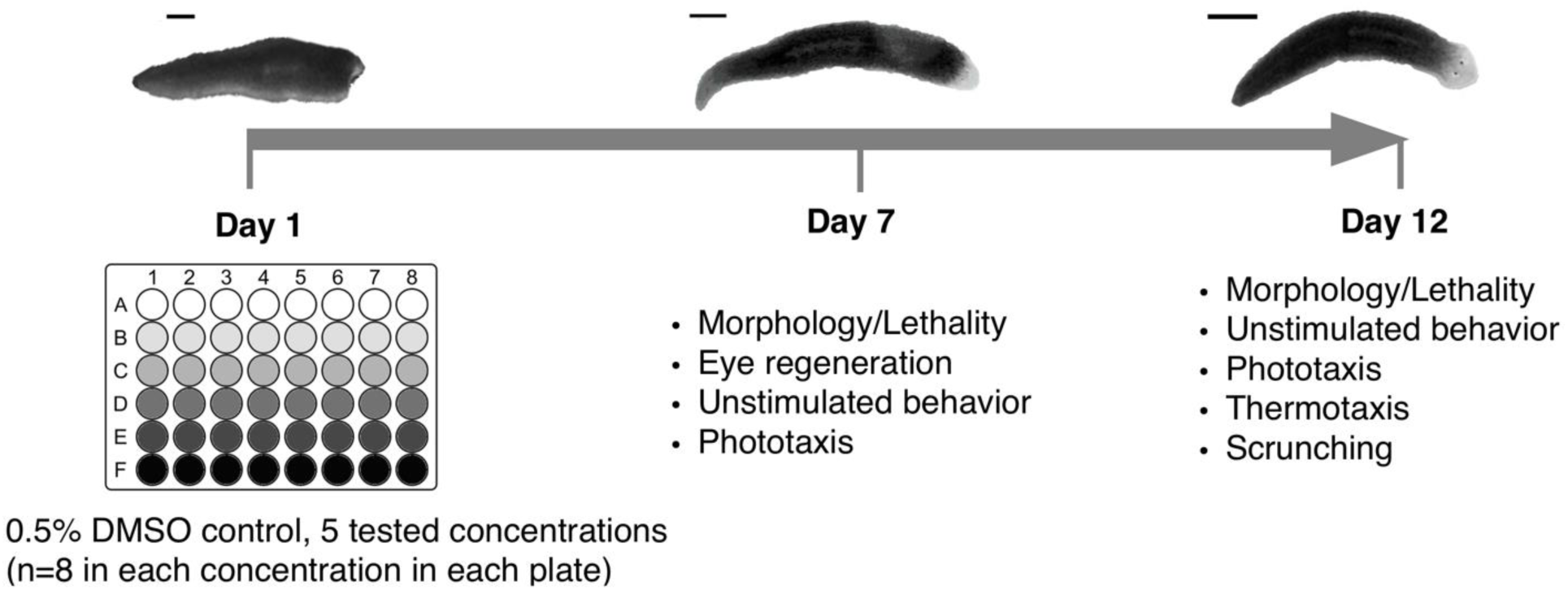
Schematic of overall screen flow in the planarian system. Exposure lasted 12 days. Amputated planarians (amputation at day 1) and adult planarians are loaded into 48-well plates (one worm per well, and one worm type per plate) with the test chemicals at day 1, and assayed for different morphological and behavioral endpoints at day 7 and day 12. Scale bar: 1mm.

A detailed description of our automated planarian screening platform and statistical workflow can be found in (Zhang et al., 2018). Briefly, adult and regenerating planarians were assessed using morphological and behavioral endpoints at day 7 and day 12 (Figure 1). Since dead worms generally disintegrate, lethality was quantified by automatic determination of the presence or absence of a body in each well. Eye regeneration was quantified by determining the percentage of planarians with 2 eyes regenerated (normal regeneration). For behavioral endpoints, we used computational image analysis to quantify animal behavior to assay unstimulated locomotion, negative phototactic behavior in response to blue light, the ability to react to a heat gradient in thermotaxis, and the ability to scrunch (musculature-driven, oscillatory escape gait (Cochet-Escartin et al., 2015)) in response to noxious heat. For statistical analysis, data from the replicate runs was compiled and each chemical concentration was compared with its own in-plate DMSO controls. Statistical significance was determined using either a one-tailed Fisher’s exact test for lethality, eye regeneration, phototaxis, and scrunching endpoints; Mann Whitney U-test for thermotaxis and unstimulated behavior; or two-tailed t-test for unstimulated behavior (depending on normality of the sample) using a significance level of 0.05. To extract biologically significant results from the statistically significant hits, “biological relevancy” cutoffs, based on the distribution of all control populations, were used to exclude hits that fell within the variability of the DMSO controls. Lastly, hits generated from inconsistent data from the different replicates were also excluded. For each endpoint, the lowest concentration which showed a significant effect (statistically and biologically) was defined as the lowest-effect-level (LEL). For each chemical, the overall LEL was determined as the most sensitive LEL when considering all endpoints. All image analyses and statistical analyses were performed in MATLAB.

The same experimental procedures and data analysis methods as described in detail in (Zhang et al., 2018) were used, except for the following differences in FR screen 2, building upon our experiences from FR screen 1: Firstly, since we found that food quality affects planarian fitness and sensitivity to chemicals (Zhang et al., 2018), we fed planarians used in FR screen 2 commercial freeze-dried organic chicken liver (Amazon, Seattle, WA) to better control food quality and thus minimize animal fitness variability. Secondly, “biological relevance cutoffs”, used to minimize false positives by accounting for variability of the solvent controls in each endpoint (Zhang et al., 2018), were updated by using the combined DMSO control samples from the previous NTP 87-compound screen (n=87 control samples) and the later FR-only screen (n=18 control samples) for a total control population of n=105 samples, with each sample consisting of n=24 planarians. Thirdly, the background noise speed threshold used in the phototaxis assay was recalculated based on the mean speed in the unstimulated behavioral assay of the expanded DMSO controls. Hits due to hyper- (rather than hypo-) activity in unstimulated behavior or hits which were concentration-independent were not included, because they were not conserved when compiling all 6 runs together, suggesting they may be artifacts.

### 2.3. Robustness analysis of planarian screening platform

The robustness of our planarian screening platform was evaluated by comparing the results obtained from six replicate runs (n=8 planarians per condition per replicate) from the two independent FR screens, FR screen 1 and 2. It is reasonable to assume that using all 6 replicates provides the most accurate results given the large sample size (n=48). We therefore compared the results of compiling different numbers of replicates (3, 4, or 5) with the results obtained by compiling 6 replicates. As described previously (Zhang et al., 2018), we used a 3-step (statistical test, biological relevance cutoff, and inconsistency check) statistical workflow to determine whether a chemical showed bioactivity, defined as a significant adverse effect, in any endpoint. Herein, we refer to the determination of bioactivity (i.e. bioactive or inactive) in each endpoint as a “readout” to distinguish it from a bioactive “hit”, which refers to having an adverse effect at an endpoint. Since the biological relevance cutoffs are meant as a means to distinguish normal wild-type noise levels, biological relevance cutoffs were determined based on the total control population of n=105 control samples (see Section 2.2), and were applied equally to these different sets of compiled data (i.e. the same cutoffs, determined from this total control population, were used for 3, 4, 5, or 6 replicates). For the inconsistency check, instances were excluded where more than half of a set of replicates (i.e. >1 out of 3 replicates, >2 out of 4 replicates, >2 out of 5 replicates, or >3 out of 6 replicates) were within the biological relevancy cutoffs. Although the majority of the experimental procedures were consistent between the two screens, some differences existed which could affect the reproducibility of the screens. For example, varying food quality (between different home-made batches and between home-made and commercial sources) affects animal health and thus affects the animals’ sensitivity to chemical exposure (Zhang et al., 2018). Different chemical batches can also cause variability due to differences in stock concentrations and potential variability in the serial dilution process. Therefore, we chose replicates (different plates from different days) with the most similar conditions (i.e. in the same screen) to minimize experimental variability and to focus on the robustness of the system itself. To compile 3 replicates, we chose the 3 replicates from FR screen 2. To compile 4 (or 5) replicates, we chose 1 (or 2) replicate(s) from FR screen 1 in addition to the 3 replicates from FR screen 2. Firstly, overall chemical bioactivity (i.e. whether the chemical was bioactive in any endpoint or inactive in all endpoints) in different sets of replicates (3, 4, 5, or 6) was compared. Secondly, concordance of the bioactivity of the readouts (i.e. whether the chemical was deemed bioactive or inactive in a specific endpoint) between the data using 3, 4, or 5 replicates and the data using 6 replicates were compared. Lastly, the concordance of the sensitivity of the concordant bioactive readouts (i.e. LEL of the bioactive readout) was compared between the data using 3, 4, or 5 replicates and the data using 6 replicates.

### 2.4. Comparison with other alternative models

To directly compare our results with published data from other alternative models, we consulted the literature (Behl et al., 2015; Behl et al., 2016; Jarema et al., 2015; Noyes et al., 2015) and focused on studies which had screened the majority of the same FRs. Studies which only tested one or a few of these FRs were not included in the comparison. We extracted the points-of-departures (PODs) and LELs reported in these papers for relevant developmental and developmental neurotoxic endpoints and focused only on the concentration-dependent hits for LELs to facilitate cross-system comparisons. Specifically, we extracted PODs for developmental toxicity and developmental neurotoxicity reported in (Behl et al., 2015), LELs for developmental behavior in (Jarema et al., 2015), LELs for larval development in (Behl et al., 2016), and LELs for all endpoints reported in (Noyes et al., 2015). Only data for regenerating planarians was used for this analysis as the comparative studies used developing models.

To quantitatively determine the agreement between the different alternative animal studies, Cohen’s kappa (κ) was calculated for each pairwise comparison where κ = (observed agreement - probability of chance agreement)/(1- probability of chance agreement) (McHugh, 2012). The 95% confidence intervals were quantified as κ ± (1.96 x the standard error). Of note, the Noyes et al. study showed a κ=0 for all comparisons because it had no inactives in this set of 9 FRs, leading to a high probability of agreement due to chance.

### 2.5. Relevance of findings in planarians to human biomonitoring data

To provide toxicologists and regulators with context of how toxicity in planarians may relate to human exposure levels, we compared the concentrations which elicited toxicity in planarians to measured or estimated human plasma concentrations of several of the tested FRs using high throughput toxicokinetic modeling (HTTK) (Rotroff et al., 2010; Sipes et al., 2017; Wetmore et al., 2012; Wetmore et al., 2013). The underlying model assumes that *in vitro* media concentrations are equivalent to human plasma concentrations and is traditionally used for cell-based or high-throughput screens. While recognizing the limitation that the planarian is an integrated system with more extensive kinetics than cell system models, this effort provides a baseline comparison that can be adjusted with increasing information of planarian kinetics. As such, this work directly compares estimated or measured (wherever available) human plasma concentrations with the regenerating planarian LEL media concentrations. Human biomonitoring data were unit-converted using HTTK modeling to estimate internal plasma concentrations in children and consisted of breast milk, handwipes, and house dust, and plasma and cord-blood serum levels of FRs as previously described (Alzualde et al., 2018). Briefly, the 3-compartment model in the HTTK R package (version 1.7) (Pearce et al., 2017; Wambaugh et al., 2015) with input chemical parameters from ADMET Predictor 7.2 (Simulations Plus, Inc., Lancaster, CA, USA) was used. Child peak plasma concentrations (over 1 year) were estimated for TPHP and TMPP from breastmilk, child hand wipe, and house dust samples (Kim et al., 2014; Sugeng et al., 2017). Adult plasma values for TBBPA and BDE47, as well as cord blood serum for TBBPA and child plasma for BDE47 were used (Cariou et al., 2008; Stapleton et al., 2012; Wang et al., 2013). Exposure values obtained were the range, as well as the geometric mean for: adult plasma, child plasma, cord serum, and TPHP child handwipe; median for: breastmilk and TMPP child handwipe, and maximum median for house dust.

### 2.6. Cholinesterase activity staining in OPFR treated planarians

To determine whether OPFR toxicity was correlated with inhibition of acetylcholinesterase activity, adult planarians were incubated in either 0.5% DMSO, 10 µM EHDP (an active OPFR in planarians) or 100 µM TCEP (an inactive OPFR in planarians) for 12 days and then fixed and stained to visualize acetylthiocholine catalysis as previously described (Hagstrom et al., 2018a; Zheng et al., 2011), with a staining incubation of 3.5 hours. The staining was performed in duplicate on separate days on a total of 6 worms per chemical (3/day) and 12 DMSO controls (6/day). Supplementary Figure 1 is representative of all imaged worms.

## 3. Results

A library of 15 FRs and 2 negative controls (acetaminophen and L-ascorbic acid) (Table 1), was screened in adult and regenerating planarians to assess their effects on mortality, eye regeneration, and behavior. These chemicals were screened twice, once as part of the NTP 87-compound library screen (FR screen 1) (Zhang et al., 2018) and once as a new, independent screen of the FR-specific subset of a newly obtained version of the library (FR screen 2), with 3 replicate runs in each screen (total n=24 per screen). Therefore, we compared the results obtained from these two independent screens using the same screening methodology to evaluate the robustness and reproducibility of our planarian screening platform. Moreover, as our previous study of the NTP 87-compound library focused on the screening methodology and major trends seen in the screen, herein we provide more in-depth insight of the toxicological profiles induced by this discrete subset of chemicals. The raw and analyzed data are available in Mendeley Data and Supplementary File 1, respectively.

### 3.1. Robustness of the planarian FR screen

To evaluate the robustness and reproducibility of our planarian screening platform, we compared the results obtained from compiling different numbers of replicates from the two independent screens (FR screen 1 and FR screen 2) (Figure 2). Compiled results for the individual replicates and different sets of replicates can be found in Supplemental File 1. Of the 18 screened chemicals, IPP and TMPP were provided by two different suppliers for the two screens (Table 1). Therefore, given potential unknown variability between the different chemical batches, IPP and TMPP were excluded from the robustness evaluation. Notably, a recent screen with developing zebrafish found significant readout differences between IPP stocks from three different suppliers (Noyes et al., 2015), confirming potential variability among suppliers.

**Figure 2.**
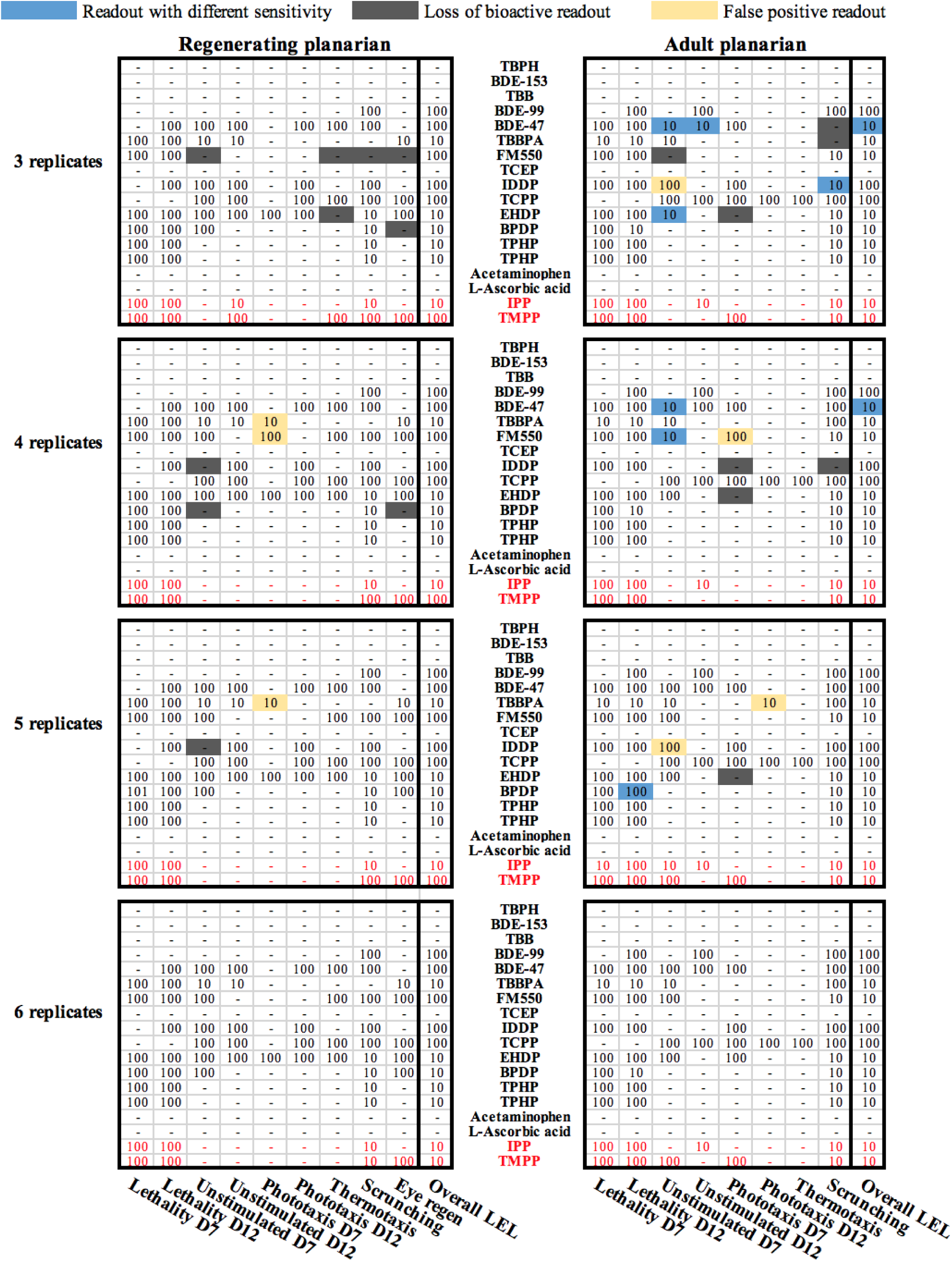
Comparison of lowest-effect-levels (LELs) identified for each endpoint by compiling data from 3, 4, 5, or 6 replicates in regenerating and adult planarians. Readouts, i.e. whether bioactivity was detected in a particular endpoint, were determined for each endpoint using 3, 4, or 5 replicates from the 2 screens and compared with the readouts determined from 6 replicates. “-” indicates the chemical was inactive in that endpoint. Loss of a bioactive readout (deemed bioactive in 6 replicates but not 3, 4, or 5) are in dark grey. False positive readouts (deemed bioactive in 3, 4, or 5 replicates but not 6) are in yellow. Endpoint readouts which differed in sensitivity/potency (still bioactive in both screens but with different LELs) are in blue. The overall LEL of each chemical is listed in a separate column. IPP and TMPP (in red text) were excluded from the robustness evaluation since the chemical supplier differed between the two screens.

To determine the robustness of our data, we evaluated both the overall agreement on bioactivity identification and potency (LELs) from any endpoints, and the agreement on bioactivity and LEL per endpoint when comparing across different numbers of compiled replicates. Firstly, we evaluated the key findings, i.e. general determination of toxicity from any endpoint and overall potency. In all sets of compiled data (3, 4, 5, or 6 replicates), the same 10/16 chemicals, including the duplicate TPHP, caused adverse effects in at least one assay endpoint in both adult and regenerating planarians. Thus, the overall chemical hit-call was unchanged regardless of the number of replicates used. Furthermore, we also found that the sensitivity of the system generally did not differ among the different numbers of replicates as the overall LELs, i.e. the most sensitive LEL from any assay endpoint for each chemical, were the same for almost all bioactive chemicals in 3/4/5 replicates versus 6 replicates. Only BDE-47 had an overall LEL, which differed depending on the number of replicates analyzed. The overall LEL for adult planarians exposed to BDE-47 was lower (i.e. more potent) when using 3 and 4 replicates compared to using 6 replicates. Significant effects were found at 10 µM BDE-47 in day 7 and 12 unstimulated behavior when analyzing 3 or 4 replicates but not 6, because in FR screen 1 BDE-47 generally showed decreased sensitivity in this assay, which we attribute to differences in animal fitness caused by differences in food quality (Zhang et al., 2018).

Secondly, because one of the strengths of our planarian screening platform is the ability to distinguish effects on different neuronal functions using various behavioral readouts, we compared the bioactivity of each endpoint readout using the compiled data of 3, 4, or 5 replicates with the compiled data of all 6 replicates (Figure 2, Table 2). Of the 144 readouts in regenerating planarians and 128 readouts in adult planarians, the bioactivity (either bioactive or inactive) determined with 6 replicates was highly concordant (≥ 96%) with that of 3, 4, and 5 replicates (Table 2). Differences were mostly due to loss of detection of bioactive readouts in 3, 4, or 5 replicates (compared to 6 replicates), although there were a few readouts which were found to be bioactive in 3, 4, or 5 but not 6 replicates. Generally, loss of bioactive readouts in 3, 4, or 5 replicates was correlated with fewer replicates, as the smaller sample size limited the ability to detect statistically significant effects. However, this primarily only occurred at concentrations which also yielded significant lethality. Thus, in these cases, there was not enough data from alive animals to yield a statistically significant effect on a morphological or behavioral endpoint. This suggests that some specific defects (e.g. scrunching) may be masked by overt systemic toxicity and lethality, especially in smaller sample sizes. The only exception to this trend was IDDP, where differences in readout bioactivity were not due to low sample size but instead from inconsistency among the different replicates from the two independent screens. Furthermore, we evaluated sensitivity concordance by comparing the LELs determined from the different replicate sets for the concordant bioactive readouts (Table 2) to determine whether the same concentration-dependent phenotypic profiles were identified in the different replicates. In regenerating tails, all concordant bioactive readouts for all endpoints had the same LELs across all sets of compiled data. In adult planarians, ≥ 97% of the concordant bioactive readouts had the same LELs using 3/4/5 replicates compared with using 6 replicates. Together, these data show the high reproducibility among the different replicates in both endpoint-specific bioactivity and potency and that our current screening strategy using 3 replicates is sufficiently robust.

**Table 2.** Bioactivity concordance of readouts and sensitivity concordance of concordant bioactive readouts of the data using 3, 4, or 5 replicates compared with the data using 6 replicates. Data are shown for regenerating (R) and adult (A) planarians. IPP and TMPP were excluded from the analysis. Bioactivity concordance of readouts was determined as the number of readouts showing the same bioactivity (bioactive or inactive) as using 6 replicates, out of the total number of readouts (9 for regenerating and 8 for adult planarians) for all 16 chemicals. Sensitivity concordance of concordant bioactive readouts using 3/4/5 replicates was determined as the number of bioactive readouts with the same LEL as in 6 replicates, out of the total number of concordant bioactive readouts (numbers in first two columns) for all 16 chemicals in this set of compiled data.

|  | **Bioactivity concordance of readouts** |  | **Sensitivity concordance of bioactive concordant readouts** |  |
| --- | --- | --- | --- | --- |
|  | R<br>(144 readouts) | A<br>(128 readouts) | R (concordant<br>bioactive readouts) | A (concordant<br>bioactive readouts) |
| **3 replicates** | 138 (96%) | 123 (96%) | 138 (100%) | 119 (97%) |
| **4 replicates** | 139 (97%) | 124 (97%) | 139 (100%) | 123 (99%) |
| **5 replicates** | 142 (99%) | 124 (97%) | 142 (100%) | 124 (100%) |

TPHP was screened as a duplicate in FR screen 2 (FR-specific library screen). The results of the duplicates were consistent in any set of compiled data, being bioactive in the same endpoints at the same LELs (Figure 2), underlining the robustness and reproducibility of our planarian screening platform.

### 3.2. OPFRS are more potent than BFRs in planarians

After confirming our data is robust and reproducible, we used the compiled data from all 6 replicates to delve into the details of FR toxicity. Eleven of the 15 unique FRs caused adverse effects in both adult and regenerating planarians in at least 1 endpoint (Figure 3). The duplicate chemical, TPHP, was consistent between batches and thus the results are only shown once. The bioactive FRs consisted of 3 of the 6 BFRs (BDE-99, BDE-47, TBBPA), 7 of the 8 OPFRs (IDDP, TCPP, EHDP, IPP, BPDP, IPP, TMPP, TPHP), and the FM550 mixture. TBPH, BDE-153, TBB, TCEP, as well as the two negative control chemicals, were inactive in both worm types at the tested concentrations. All chemicals were screened at 5 test concentrations (see Materials and Methods, Section 2.1); however, hits were only observed at the two highest concentrations, 10 and 100 µM. Due to poor solubility, BDE-153 was tested at a maximum concentration of 50µM, which may explain its inactivity, since the other BDE compounds were only active at 100µM.

**Figure 3.**
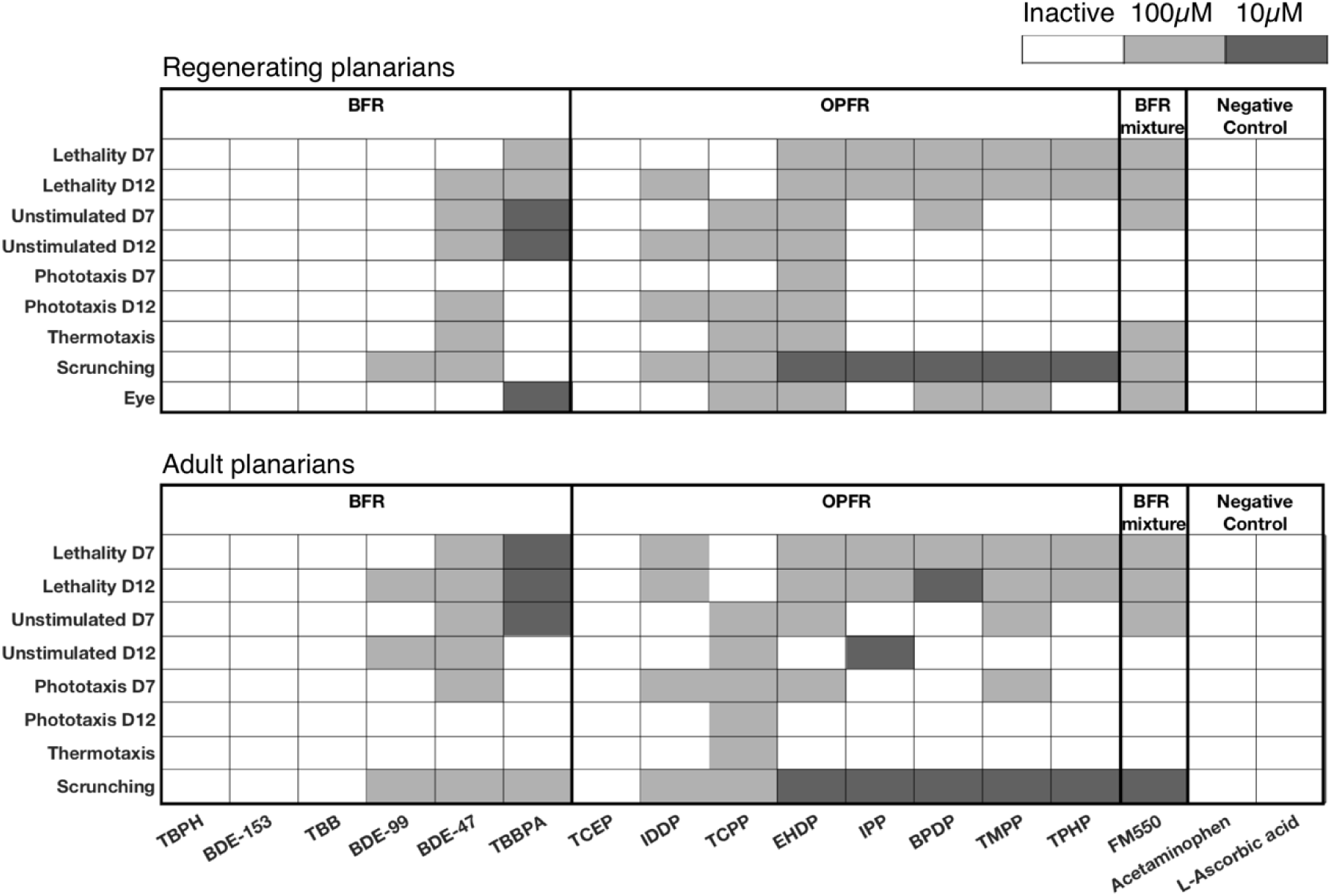
Overview of the planarian screening data of 15 unique flame retardants and 2 negative controls using 6 replicates. Heat maps of effects of brominated flame retardants (BFRs), organophosphorus flame retardants (OPFRs), a BFR mixture, and negative controls on regenerating (Top) and adult (Bottom) planarians for all endpoints with lowest-effect-level shaded. Only the highest two of the five tested concentrations showed effects in planarians. TPHP was screened as a duplicate in FR screen 2, but only shown once here since the results were consistent. Note that BDE-153 was screened at a maximum concentration of 50 µM.

Using the high-content data obtained by screening for effects on planarian morphology and a variety of diverse behaviors, we determined whether any specific toxicities were induced by the different FRs. Eight FRs in regenerating planarians (BDE-99, TBBPA, TCPP, EHDP, IPP, BPDP, TMPP, and TPHP) and 6 FRs in adult planarians (TCPP, EHDP, IPP, BPDP, TMPP, and TPHP) caused adverse outcomes on morphological and/or behavioral endpoints at concentrations where animal viability was not affected, suggesting specific non-systemic toxicity at these concentrations. Although causing sublethal effects in regenerating planarians, two BFRs, BDE-99 and TBBPA, only showed toxicity concomitant with lethality in adult planarians, due to increased sensitivity to lethality in the adult worms. Similar increased sensitivity of adult planarians to the lethal effects of some chemicals has been observed previously (Hagstrom et al., 2015; Zhang et al., 2018).

Next, we further examined which specific endpoints were affected by the sublethal concentrations of the different FRs. Scrunching, a planarian musculature-driven gait used as an escape response to specific adverse stimuli such as noxious heat (Cochet-Escartin et al., 2015), was the most sensitive endpoint with 7 and 6 FRs causing scrunching defects at sublethal concentrations in regenerating and adult planarians, respectively. To a lesser extent, sublethal effects were also seen on unstimulated behavior (regenerating planarians in TBBPA and adult planarians in IPP) and eye regeneration (TBBPA and TCPP). Interestingly, in addition to effects on eye regeneration, TCPP affected almost all behavioral endpoints (with the exception of day 7 phototaxis in regenerating planarians) in both worm types in the absence of lethality. Together, these toxicological profiles demonstrate the power of the large number of diverse endpoints to discern specific toxic effects from systemic toxicity and pinpoint effects on specific neuronal functions, such as neuromuscular communication.

Our planarian HTS platform has the unique advantage of allowing direct comparisons of chemical effects on adult and developing/regenerating animals. By doing so, we found TBBPA caused a developmental selective defect on day 12 unstimulated behavior, since effects on this endpoint were seen at 10µM in regenerating but not adult planarians (Figure 3). Of note, this concentration did induce lethality in 24% of adult planarians at day 12 (see Supplemental File 1). However, this low level of lethality, although statistically significant, may suggest that any overt systemic toxicity at this concentration would not be potent enough to mask effects in other endpoints. Furthermore, TBBPA and TCPP caused eye regeneration defects at a sublethal concentration. Thus, while most compounds adversely affected both adult and developing animals, some compounds displayed additional developmental-specific toxicities.

Lastly, to ask whether OPFRs are safer alternatives to BFRs, we compared the toxicological profiles induced by the two types of FRs. The OPFRs generally showed higher potency than the BFRs with LELs of 10µM in 5 out of 7 bioactive OPFRs, whereas the bioactive BFRs, except for TBBPA, had LELs of 100µM. TBBPA had a LEL of 10µM in both adult and regenerating planarians, with the most endpoints affected at this concentration, compared to the other FRs.

Taken together, the bioactive OPFRs were at least as potent as BDE-47, which was phased-out due to concerns about its toxicity (Stapleton et al., 2014), emphasizing the importance of evaluating the toxicity of these supposedly safer replacement FRs. Almost all bioactive OPFRs (6/7 and 5/6 for regenerating and adult planarians, respectively) induced specific sublethal effects on scrunching. This was not seen to the same extent in the BFRs, where only 1 bioactive BFR (BDE-99) showed sublethal effects on scrunching and only in regenerating planarians. Thus, by assaying a variety of different neuronal functions in planarians, we were able to differentiate toxicities elicited by different chemical domains, suggesting that our system provides specificity.

### 3.3. Concordance of bioactive hits between planarians and other alternative models

While several of the 15 FRs that we screened have been previously studied in other systems, including rodents (Cope et al., 2015; EFSA, 2011; Moser et al., 2015; Nakajima et al., 2009; National Toxicology Program, 1991; Viberg and Eriksson, 2011) and other alternative models (Alzualde et al., 2018; Bailey and Levin, 2015; Cano-Sancho et al., 2017; Glazer et al., 2018; Oliveri et al., 2015; Slotkin et al., 2017; Usenko et al., 2016), only a handful of studies have performed direct multi-FR comparisons as provided here. Fourteen of the 15 unique FRs tested in this screen have been previously studied in developing zebrafish, nematodes, and *in vitro* cell-based (mouse embryonic stem cell, human neural stem cell, and rat neuron) systems with some FRs showing developmental toxicity or DNT. Thus, herein, we compared our planarian data with these published results (Behl et al., 2015; Behl et al., 2016; Jarema et al., 2015; Noyes et al., 2015) to contextualize our findings (Table 3, Figure 4). The FM550 mixture was excluded from this comparison because it was not screened in any of these previous studies. Comparisons were made with the overall regenerating planarian LEL, i.e. most sensitive LEL from any endpoint, determined from the compiled data of 6 replicates from FR screens 1 and 2 (see Methods Section 2.4, Table 3 and Figure 4).

**Table 3.**
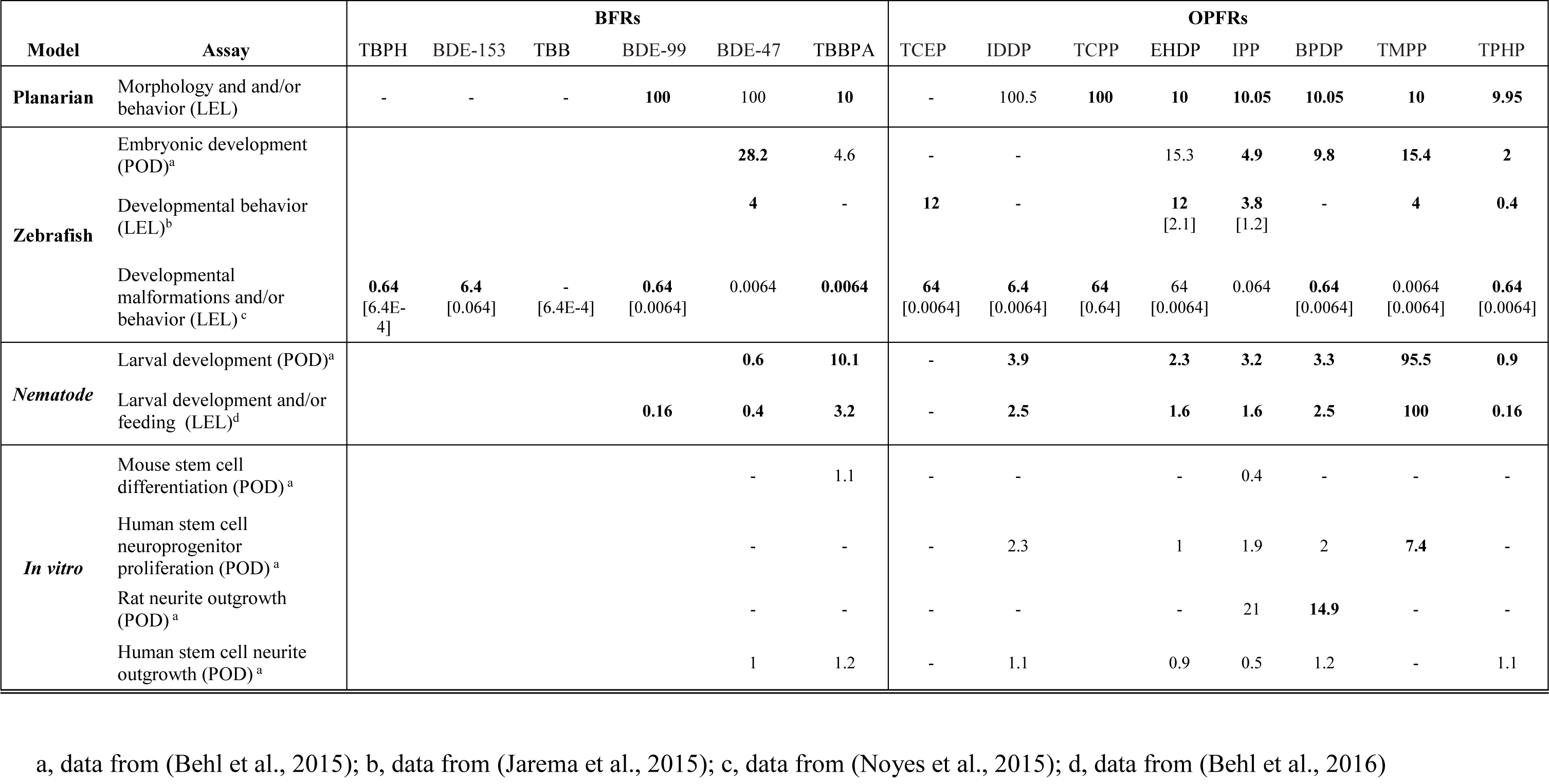
Comparison of developmental and/or developmentally neurotoxic endpoints used to evaluate FR toxicity between regenerating planarians and different alternative developing models. Lowest-effect-level (LEL) or point-of-departure (POD) for each model are listed in µM. Dash (-) indicates that there was no toxicity detected at the tested concentrations. An empty cell indicates that this chemical was not tested in this model. The number in brackets represents the LELs when considering concentration-independent and hyper-active effects. Bold text indicates that the effects were detected at sublethal concentrations.

**Figure 4.**
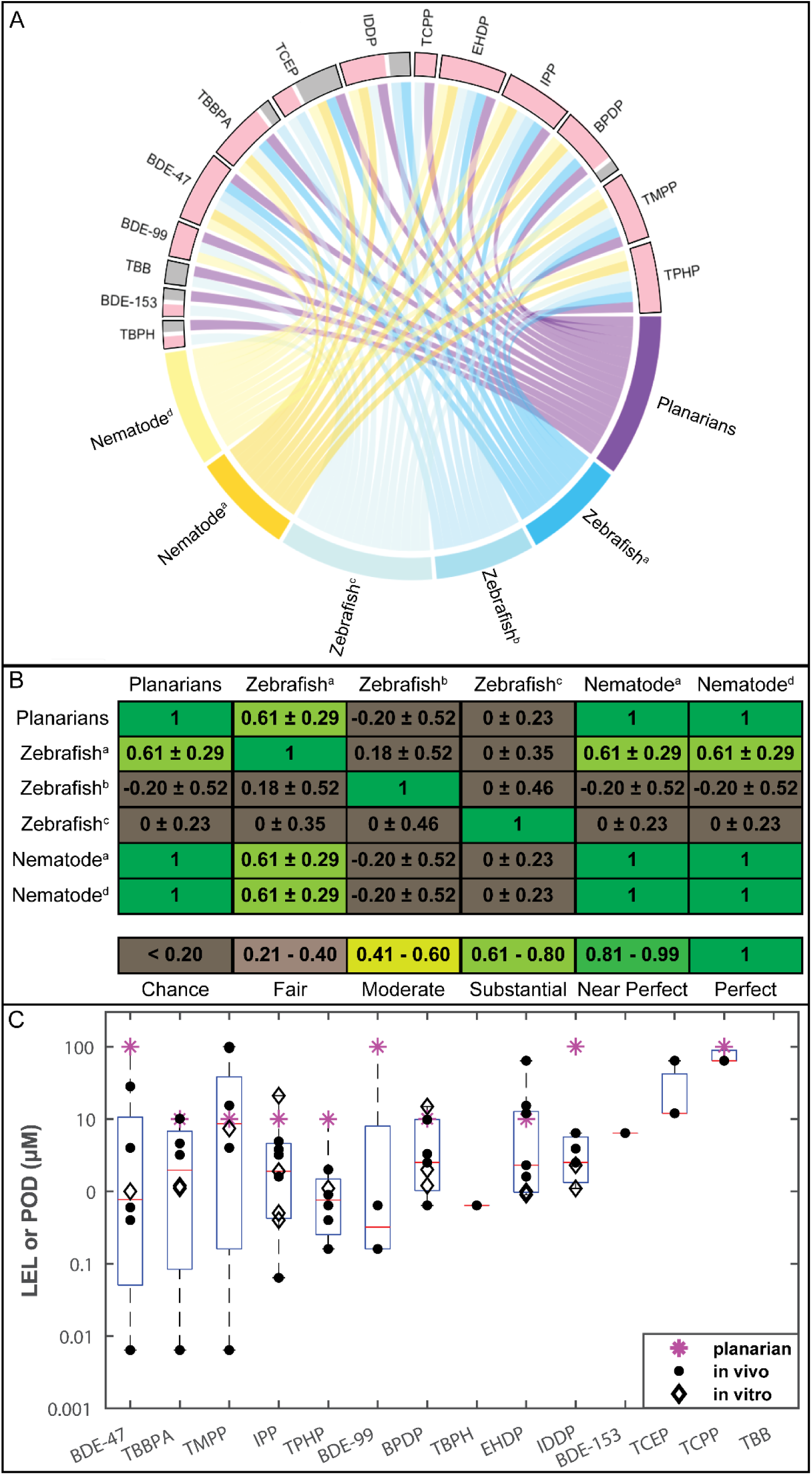
A. Interaction of bioactivity of the 14 flame retardants (FRs) with the different tested alternative animal models. Each box represents the interactions of this FR with the different models and studies, coded by different colors, where active interactions are in pink and inactive interactions are in grey. FRs with both active and inactive interactions signify that the chemical showed different effects in different studies/models. B. Quantification of concordance between alternative animal models for 9 FRs screened in all alternative animal models using Cohen’s kappa (κ), listed as κ ± 95% confidence intervals (McHugh, 2012). Of note, the Noyes et al study showed a kappa of 0 for all comparisons because it had no inactives in this set of 9 FRs. For (A) and (B), studies are listed as: a, data from (Behl et al., 2015); b, data from (Jarema et al., 2015); c, data from (Noyes et al., 2015); d, data from (Behl et al., 2016). C. Comparison of toxicity potency of 14 FRs in regenerating planarians and other alternative developing models. The data for the individual studies are overlaid on a box and whisker plot compiling the lowest-effect-levels (LELs) and points-of-departure (PODs) for the different FRs in all the different models. Stars represent planarian LELs, filled dots represent LELs/PODs in developing zebrafish and nematode models (Behl et al., 2015; Behl et al., 2016; Jarema et al., 2015; Noyes et al., 2015), and diamonds represent PODs in *in vitro* cell-based models (Behl et al., 2015). The x-axis shows the FRs in decreasing order of potency (lowest effect concentration).

First, we compared the overall chemical hit calls between the different systems to determine whether the same FRs were identified as bioactive, i.e. caused significant adverse effects, in the different models. Thirteen of the 14 shared FRs were previously found to be bioactive in at least one model; however, not all 14 FRs were tested in all systems or studies. IPP was the only FR found to be bioactive in all tested systems. TBB was the only FR which was inactive in all screens; however, it was only tested in our study and the (Noyes et al., 2015) zebrafish study. Generally, the alternative animal models detected more bioactive FRs than the *in vitro* models (Table 3). Of the 9 FRs tested in the *in vitro* models, 8 were bioactive in any model, with at most 7 detected in any one model (human stem cell neurite outgrowth). The majority of effects caused by the FRs in the *in vitro* models occurred concomitantly with cytotoxicity, suggesting non-specific effects.

Due to the increased specificity and higher similarity to our planarian screening platform, we focused our concordance analysis on the alternative developing/regenerating animal models (planarians, nematodes, and zebrafish). The chemical hit calls for the 14 FRs tested in these models were highly concordant. The two invertebrate systems, nematodes and planarians, had perfect hit concordance for the 10 FRs tested in both systems. These 10 FRs were also tested in at least one zebrafish study and showed high concordance among the three models (considering all studies together) with the same 9 chemicals eliciting adverse effects in at least one study from each system (Table 3, Figure 4A). Of these 10 FRs, only TCEP had discordant results between the different systems, as it was bioactive in two of the zebrafish studies, but inactive in the third zebrafish study and in nematodes and planarians.

As this example illustrates, some discrepancies were found across different studies using the same model. Detailed comparisons are shown in Table 3 and Figure 4A. We used Cohen’s kappa, which also considers agreement due to chance (McHugh, 2012), to quantify concordance across the different studies for the 9 shared FRs (Figure 4B). As expected, we found perfect agreement between the two nematode studies and our planarian study. However, all other comparisons yielded substantial agreement or only agreement due to chance, even when comparing different studies in the same system, demonstrating that methodological differences between labs studying the same system can be as impactful as interspecies differences. This emphasizes the importance of transparency, so results from different studies can be adequately compared and compiled to leverage the strength of multi-system testing.

We also compared the sensitivity (i.e. chemical potency) of our regenerating planarian screening data to those of the comparative alternative model studies (Figure 4C, Table 3). Notably, some of these studies, including ours herein, evaluated toxicity potency using the nominal test concentrations (i.e. LEL) (Behl et al., 2016; Jarema et al., 2015; Noyes et al., 2015) while others used modeling approaches to calculate a point-of-departure (POD) (Behl et al., 2015). Eight of the 10 FRs causing adverse effects in planarians (BDE-47, TBBPA, BPDP, EHDP, IPP, TMPP, TPHP, and TCPP) had overall LELs (from any endpoint) in the regenerating planarian system within an order of magnitude of the effect concentration (LEL or POD) in at least one of the other studies using zebrafish, nematode, and *in vitro* cell-based models (Figure 4C, Table 3).

## 4. Discussion

### 4.1. Robustness of planarian screening platform

Robustness of screening data is indispensable for building confidence in a new testing platform. In previous proof-of-concept work (Hagstrom et al., 2015), we used post-hoc statistical power analysis to determine that n=24 specimen per condition were necessary to achieve sufficient power. Here, we experimentally verified the validity of our current test strategy, testing in 3 replicate runs with n=8 each, through direct comparison of the reproducibility of data obtained from using 3 (n=24), 4 (n=32), or 5 replicates (n=40) with 6 replicates (n=48). We found that our planarian platform is robust and that 3 replicates were sufficient to identify all chemical hit-calls, distinguishing the same bioactive chemicals as 6 replicates. Moreover, readout bioactivity and sensitivity (i.e. LELs) of the different endpoints were highly concordant among the different sets of replicates. Loss of bioactive readouts with less replicates was only found for morphological or behavioral endpoints at lethal concentrations. Since all endpoints, except lethality, were quantified from the data from living animals, high lethality led to a small sample size and limited the power to detect a statistically significant effect. Moreover, lethality suggests overt systemic toxicity at these concentrations which may have masked effects at more specific endpoint readouts. Given the overt toxicity at lethal concentrations, many labs only evaluate more sensitive morphological or behavioral effects at sublethal concentrations, disregarding effects at lethal concentrations (Alzualde et al., 2018; Jarema et al., 2015; Truong et al., 2014). All together, these data indicate that different number of replicates (from 3-6) yield very similar results, validating that our current screening strategy using 3 replicates is able to robustly identify chemical bioactivity and different toxicities.

### 4.2. The role of in vitro HTS for DNT

As laid out in the introduction, few mammalian guideline studies have been conducted to evaluate the DNT of chemicals, including the FRs tested herein. High-throughput *in vitro* assays, coupled with bioinformatic analysis, have emerged to fill this gap by quickly and efficiently evaluating potential chemical toxicity. However, our comparison of several human and rodent cell culture assays with non-mammalian whole animal models found that the alternative animal models were better at identifying specific developmental or developmental neurotoxic effects at sublethal concentrations (Table 3). When considering all compared *in vitro* assays, 8/9 FRs were found to be bioactive; however, 6 of these bioactive FRs were only identified in 1 or 2 assays (primarily the human stem cell systems). The two rodent cell culture assays only yielded 2 hits each, both of which were also detected by the human stem cell assays. The majority of the *in vitro* effects were found concomitantly with cytotoxicity, while selective sublethal effects were more apparent in the alternative animal models. A recent study evaluating a small number of BFRs and OPFRs in rat neural stem cells and neuronotypic PC12 cells was able to distinguish effects on neurodifferentiation from cytotoxicity (Slotkin et al., 2017), suggesting that certain types of cell culture assays may be more suited for DNT studies than others. Thus, while current *in vitro* techniques are extremely powerful for rapid screening of toxicants on other organ systems, they may have limited value for detecting potential developmental toxicity or DNT, because they inherently lack the necessary complexity for detecting functional perturbations that are independent of systemic toxicity. Alternative whole animal systems present an opportunity to fill this gap between *in vitro* and mammalian animal models.

### 4.3. Need for transparency in screening methodology and analysis

As shown in Figure 4B, intraspecies differences in chemical hit-calls between different studies were as big as interspecies differences. Additionally, even when a chemical was identified as bioactive in two studies from the same model, sensitivity could vary dramatically. For example, the zebrafish data from (Noyes et al., 2015) shows increased sensitivity (up to 3 orders of magnitude difference) to 4 FRs (TBBPA, BPDP, IDDP, and IPP) when compared to the other published zebrafish studies (Table 3).

Important methodological differences in experimental design and data analysis may explain these differences. For example, we tested the FRs in log-steps, whereas several of the other studies used smaller concentration steps, thus the large dose gaps in our study limits precise comparisons of potency. In addition, how significant effects are determined can vary dramatically between laboratories. Some use statistics alone, while others, like ourselves incorporate additional stringency tests including biological relevancy cutoffs and inconsistency checks (Zhang et al., 2018). Our statistical pipeline is based on empirical observations to reduce the false positive rate, but may limit sensitivity. For example, we found our biological relevancy cutoffs removed 72% of statistically significant hits, which changed the LELs of 8 chemicals (Supplementary Table 1). However, the majority of the removed hits were concentration-independent. i.e. effects were seen at lower but not the subsequent higher concentrations, and some were hits in negative controls, together suggesting that these excluded hits are likely false positives (Zhang et al., 2018). Variability can also arise due to using LELs vs PODs to determine potency. LELs are limited to the tested concentrations, and thus the respective POD could lie somewhere within the range of the LEL and the next lower tested concentration level. Therefore, because the different experimental approaches and statistical criteria used to evaluate bioactivity and potency are as important as differences between species, transparency of methodology is crucial to make system comparisons meaningful and build confidence in these new approaches for complementing mammalian toxicity testing.

### 4.4. Comparisons of different models emphasizes the value of a battery approach

To contextualize the results of our planarian screen with what is known for FR toxicity, we compared the available data from alternative, non-mammalian animal models and found that our results were highly concordant. Eight of 10 common FRs tested in planarians, zebrafish, and nematodes were bioactive in at least one study in all three systems (Table 3 and Figure 4A). Thus, these shared bioactive FRs affected multiple models and multiple endpoints, spanning effects on development, neurodevelopment, and general neuroactivity, indicating multiple mechanisms may be involved in their toxicity. Battery approaches such as these are powerful because they allow parallel interrogation of different species and endpoints, from the cellular to organismal level, to integrate multiple lines of evidence and build confidence in preliminary hazard assessment.

In addition to comparing with other toxicological models, comparing with human environmental exposure can contextualize the relevancy of observed toxicity in a given system. Therefore, we overlaid our regenerating planarian screening results to measured or estimated *in vivo* plasma concentrations using the HTTK R-package (Figure 5) (Alzualde et al., 2018; Pearce et al., 2017). The nominal water concentrations at which effects were seen in regenerating planarians are significantly greater (1-2 orders of magnitude) than the highest estimated concentrations based on reported human exposure levels. These human exposure values have also been recently compared with the nominal and internal LELs in a developing zebrafish screen (Alzualde et al., 2018). While the zebrafish nominal LELs were generally closer to the human exposure levels than planarian nominal LELs, the majority were also outside of the range of human exposure. One possible interpretation of these data is that the toxicity observed in these alternative models are irrelevant because it occurs at higher concentrations than what was estimated from average human exposure data. However, care has to be taken when performing these comparisons, because the modeled data does not take into account differences in uptake routes and levels, metabolism, or possible tissue accumulation over time as it is modeling plasma concentration. Accumulation represents a tangible risk factor particularly for FRs as both BFRs and OPFRs have been suggested to bioaccumulate (Hale et al., 2003; Hendriks and Westerink, 2015; Hou et al., 2016; Nakajima et al., 2009). In fact, when comparing internal and nominal concentrations, a zebrafish study found vast differences among the different FRs, with up to ∼600X increases in internal FR concentrations in day 4 zebrafish larvae compared to the nominal concentrations (Alzualde et al., 2018).

**Figure 5.**
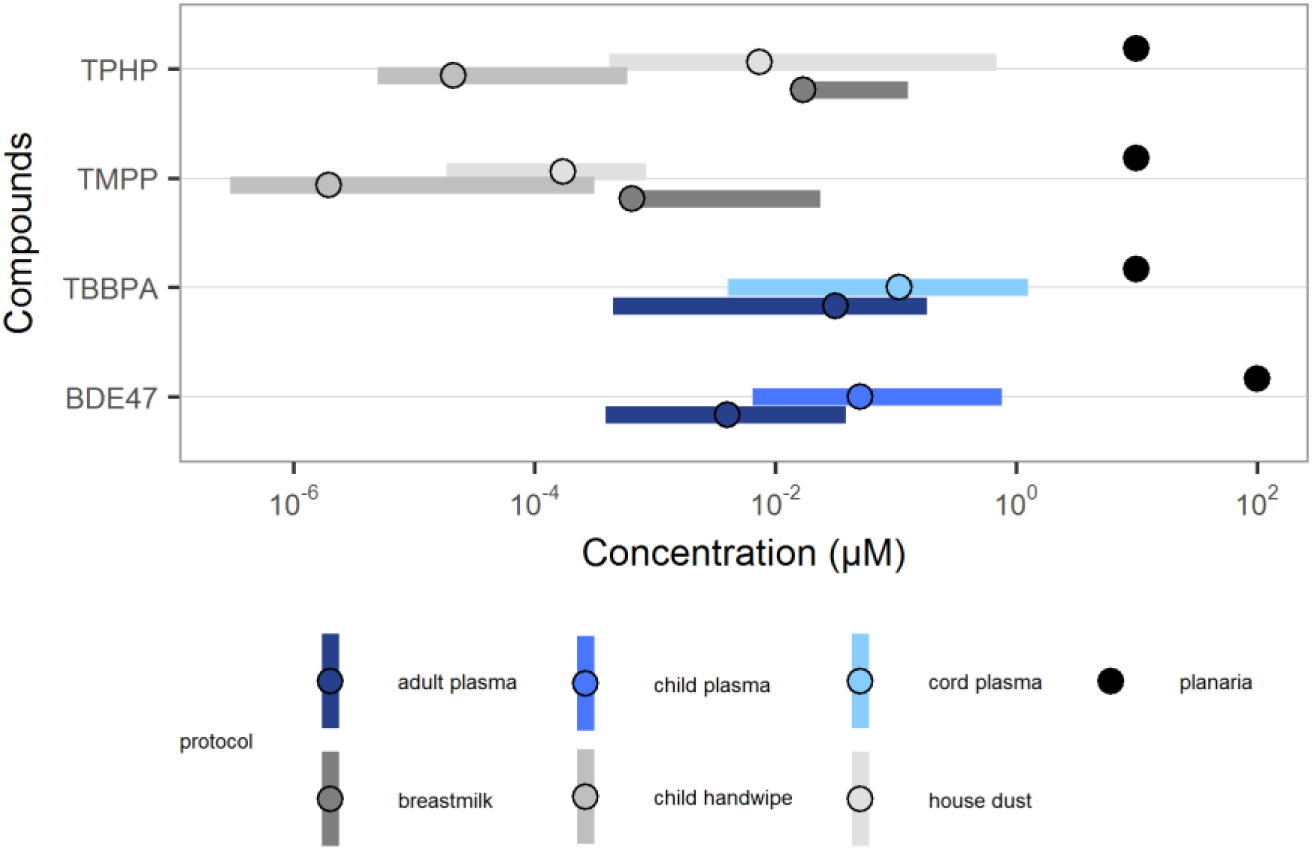
Overlay of the LELs in regenerating planarians with available human exposure data. Unit-converted measured flame retardant concentrations in adult plasma, child plasma, and cord plasma, as well as HTTK R-package estimated child plasma concentrations from breastmilk, child handwipe, and house dust were plotted (colored bars and circles). The bars represent the range, while the respective circles represent the geometric mean, median, or maximum median as noted in the Methods (Section 2.5). The lower and upper range of the unit-converted (plasma) and HTTK-estimated internal concentrations were calculated using the lowest and highest reported values, respectively. The LELs (nominal concentrations) in planarians (black dots) were based on the most potent effect at any endpoint in regenerating planarians (Table 3 and Figure 3).

For all systems comparisons, it is imperative to understand the differences between nominal water concentrations and internal doses, particularly at the site of action. For example, although the human exposure data is based on human plasma concentrations, these may not reflect the relevant concentrations at the site of action, i.e. the nervous system. Similarly, the toxicity in the alternative animal models discussed herein are based on nominal water/media concentrations and not doses. Since the effective internal concentrations in the animals are unknown, strong conclusions of comparative sensitivity and their relevancy to human exposure are difficult to make. Thus, understanding the potential for toxicity, even at nominal concentrations which may, at first glance, seem irrelevant to human exposure, are important for risk assessment.

### 4.5. Replacement FRs show comparable neurotoxicity to phased out PBDEs

In this screen, we evaluated the neurotoxicity and DNT of 15 FRs (6 BFRs, 8 OPFRs, and 1 BFR mixture) and found that 11 (3 BFRs, 7 OPFRs, and the BFR mixture) caused adverse effects in both adult and regenerating planarians. It should be noted that the 3 inactive BFRs, BDE-153, TBPH, and TBB have relatively high molecular weights (550-706 g/mol). Moreover, TBPH and BDE-153 had solubility issues at the highest tested concentration (100 and 50 µM, respectively) (Zhang et al., 2018). Thus, these issues could have contributed to their inactivity in our planarian screen. TCEP, the only OPFR inactive in our planarian screen, was also inactive in all studies (*in vivo* and *in vitro*) except for 2 zebrafish studies (Table 3) (Jarema et al., 2015; Noyes et al., 2015). Additionally, inactivity in nematodes was likely not an issue of limited test concentrations as TCEP was tested at up to 1mM without any observed effects. Only the Jarema et al and Noyes et al studies found TCEP to be bioactive in zebrafish developmental behavioral endpoints, albeit at relatively higher concentrations than the other tested FRs, suggesting relatively low potency. In agreement with the results in worms, studies with Long-Evans rats found no significant DNT was induced by oral exposure (gavage) of up to 90 mg/kg/day TCEP from gestational day 10 to weaning (Moser et al., 2015).

Several currently used FRs, including one in-use BFR, were found to be similarly or more toxic than the phased-out PBDEs in planarians. TBBPA, a BFR which is still currently widely in use, was found to cause specific, sublethal effects on unstimulated behavior (day 7 and 12) and eye regeneration in regenerating planarians (Figure 3), and was more potent than the phased-out BDE-47 and −99. In addition, effects on day 12 unstimulated behavior were selective to regenerating planarians. Together, these results suggest TBBPA caused selective adverse effects on planarian development. Although thus far guideline animal studies have not demonstrated any teratogenic effects of TBBPA (EFSA, 2011), developmental defects were observed in several alternative models compared here, including planarians, zebrafish (Noyes et al., 2015), and nematodes (Behl et al., 2015; Behl et al., 2016) at concentrations of 10 µM and below, suggesting conserved mechanisms of toxicity which may need to be re-examined in the context of human risk assessment. In addition, in mice, TBBPA has been reported to cause neurobehavioral defects after acute exposure and act as a thyroid endocrine disruptor (Nakajima et al., 2009; Viberg and Eriksson, 2011).

Similarly, almost all of the tested OPFRs, which were marketed as safer alternatives to BFRs, were found to be at least as toxic as the phased-out BDEs in planarians. Our results resonate with previous studies in nematode, zebrafish and human, mouse, and rat cell culture, which similarly found that most OPFRs had comparable activity to BFRs (Alzualde et al., 2018; Behl et al., 2015; Behl et al., 2016; Jarema et al., 2015; Noyes et al., 2015). However, because of the range of behaviors tested in our planarian platform, which assay various neuronal functions, we were able to differentiate the toxicities of OPFRs compared to the BFRs. OPFRs generally caused more sublethal, specific neurotoxic effects whereas BFRs were primarily systemically toxic, as discussed in detail in Section 4.6.

### 4.6. Behavioral profiling in planarians adds screening specificity

The majority (5/7) of the bioactive OPFRs in regenerating and adult planarians showed specific, sublethal effects on scrunching, which were not seen to the same extent in the BFRs. Scrunching is an oscillatory, musculature-driven escape gait in planarians that is induced by specific adverse stimuli such as noxious heat (Cochet-Escartin et al., 2015), but is not a generic response to all noxious environments (Cochet-Escartin et al., 2016). Although scrunching shares qualitative features, such as muscle-based motion and increased mucus secretion, with other muscle-driven body shape changes, such as peristalsis, it can be quantitatively distinguished using four characteristic parameters: frequency and asymmetry of body length oscillations, speed, and maximum elongation (Cochet-Escartin et al., 2015). Scrunching is also distinctively different from other body undulations described in the literature, including snake-like behaviors and seizures (Ross et al., 2018).

Scrunching defects, as tested in our HTS platform, therefore point to specific adverse effects on one or multiple molecular targets controlling the underlying signaling cascade from stimulus sensation to muscle contraction, mucus secretion, and locomotion. Importantly, some of the molecular targets that mediate scrunching have been previously identified. For instance, it has been shown that the Transient Receptor Potential Channel Ankyrin 1 (TRPA1) is required for proper scrunching in response to tail amputation (Arenas et al., 2017). TRPA1 is also an indirect sensor of noxious heat in planarians (Arenas et al., 2017) and we have recently confirmed that TRPA1 activation is required for heat-induced scrunching (unpublished results). Additionally, cholinergic neurotransmission, which is responsible for coordinating planarian motor functions (Nishimura et al., 2012), is likely important for scrunching, as this gait relies on muscle contraction. We have previously characterized two putative genes responsible for cholinesterase activity in *D. japonica* and found them to be sensitive to inhibition by organophosphorus pesticides and linked to behavioral defects (Hagstrom et al., 2017; Hagstrom et al., 2018a). Organophosphorus compounds share the ability to inhibit cholinesterase, though their potency can vary due to structural and pharmacokinetic differences (Taylor, 2018). Because comparable concentrations of organophosphorus pesticides, such as parathion and chlorpyrifos (Zhang et al., 2018), also cause scrunching defects, this phenotype is likely representative of the shared cholinergic toxicity. Consistent with this idea, we found an almost complete loss of cholinesterase activity in day 12 adult planarians exposed to 10 µM EHDP, which causes scrunching defects. In contrast, cholinesterase activity in day 12 adult planarians exposed to 100 µM TCEP, which was the only inactive OFPR in our planarian screen, was indistinguishable from controls (Supplementary Figure 1). Similarly, it was previously shown that DNT of 4 OPFRs tested in medaka was correlated with changes in acetylcholinesterase activity. For example, TPHP caused developmental and behavioral defects concomitant with significant acetylcholinesterase inhibition whereas TCEP, which had no significant effect on acetylcholinesterase activity, had much milder toxic effects (Sun et al., 2016b). Lack of significant effects of TCEP on acetylcholinesterase activity or transcription were also found in developing zebrafish (Sun et al., 2016a). Together, these studies indicate that the inactivity of TCEP in these different models may result from its inability to significantly inhibit acetylcholinesterase at the tested concentrations.

Our behavior-rich planarian system can be a powerful complement to existing developing zebrafish and nematode models, with their respective morphological breadth and genetic tractability. Because we can directly compare toxicities in regenerating and adult planarians, we are able to distinguish toxic from developmental toxic effects, which is not directly possible in these other systems. For example, we found developmentally selective effects on day 12 unstimulated behavior following exposure to 10µM TBBPA in regenerating but not adult planarians, suggesting TBBPA may act on specific developmental processes or on processes which are more sensitive during development. The shared effects and potencies of the other bioactive FRs on both regenerating and adult planarians suggest a potential shared mechanism of toxicity, with effects on overall neural function rather than neurodevelopment directly. Together, these examples illustrate that, in a single screen, the planarian platform is uniquely able to differentiate DNT from neurotoxicity and distinguish effects on specific neuronal functions.

## 5. Conclusions

Here, we showed, using the concrete example of screening 15 FRs, that our planarian HTS platform generates robust data and allows us to distinguish systemic toxicity from neurotoxicity and developmentally selective effects from general neurotoxicity. Additionally, the diverse behaviors tested in this planarian system allowed us to differentiate class-dictated phenotypic signatures, particularly evidenced by the shared effect of OPFRs, but not BFRs, on planarian scrunching. Concordance between this planarian study and previous studies in cell culture, nematodes and developing zebrafish, demonstrating the comparable toxicity of replacement OPFRs with BFRs emphasizes the urgent need for further evaluation of OPFRs in mammalian systems.

## 6. Supplementary data description

Compiled data for each endpoint and comparisons between individual replicates can be found in Supplementary File 1. Additional data on the differences between our determined hits and initial statistically significant effects can be found in Supplementary Table 1. Results from the cholinesterase activity staining of EHDP and TCEP can be found in Supplementary Figure 1.

## 8. Acknowledgements

The authors would like National Toxicology Program (NTP) for providing the chemical library, Andrew Hyunh and Yingtian He for help with data compilation, and Christina Rabeler for the acetylcholinesterase activity staining.

