## Supplementary Materials for "Screening for neurotoxic potential of 15 flame retardants using freshwater planarians"

**Supplementary Table 1**. Lowest effect levels (LELs) in µM for each chemical before and after applying the biological relevancy cutoffs and inter-plate inconsistency checks in adult and regenerating planarians. The originial LELs were determined based on the statistically significant hits at each endpoint. The final LELs were based on the statistically and biologically significant hits (with biological relevance cutoffs and inconsistency check applied; see Methods and Zhang et al., 2018). The dash "-" represents the chemical was inactive. Final LELs which differed from the original LELs are shaded. Supplementary File 1 includes the details of how each individual hit was affected by either biologically relevancy cutoffs or the inter-plate inconsistency check.

| | Adult planarian | | Regenerating planarian | |
|---|---|---|---|---|
| **Chemical** | **Original LEL** | **Final LEL** | **Original LEL** | **Final LEL** |
| **TBPH** | 0.01 | - | - | - |
| **BDE 153** | - | - | 0.005 | - |
| **TBB** | 0.01 | - | 0.01 | - |
| **BDE-99** | 10 | 100 | 0.01 | 100 |
| **BDE-47** | 0.01 | 100 | 0.01 | 100 |
| **TBBPA** | 0.01 | 10 | 1 | 10 |
| **TCEP** | - | - | 0.1 | - |
| **IDDP** | 0.1 | 100 | 1 | 100 |
| **TCPP** | 100 | 100 | 100 | 100 |
| **EHDP** | 1 | 10 | 0.01 | 10 |
| **IPP** | 10 | 10 | 0.1 | 10 |
| **BPDP** | 10 | 10 | 1 | 10 |
| **TMPP** | 10 | 10 | 0.01 | 10 |
| **TPHP** | 10 | 10 | 0.1 | 10 |
| **TPHP duplicate** | 0.1 | 10 | 0.1 | 10 |
| **FM550** | 1 | 10 | 1 | 100 |
| **Acetaminophen (4-hydroxyacetanilide)** | 0.01 | - | 0.1 | - |
| **L-Ascorbic acid** | - | - | 1 | - |

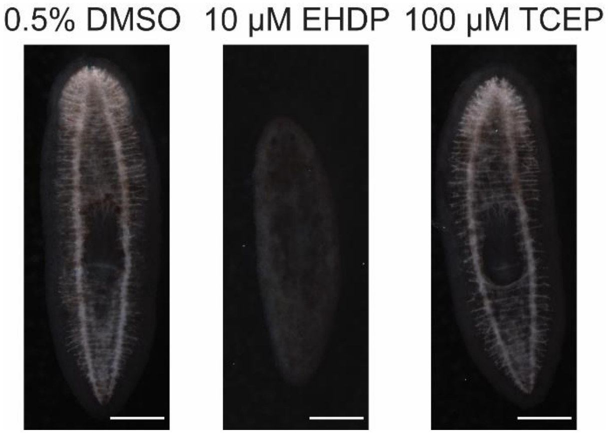


**Supplementary Figure 1**. Significant loss of cholinesterase activity in planarians exposed to EHDP but not TCEP. Representative images are shown of adult planarians incubated in the respective chemical concentrations for 12 days and then fixed and stained to visualize catalysis of acetylthiocholine (Hagstrom et al., 2017; Hagstrom et al., 2018; Zheng et al., 2011). Note that almost no activity, seen as the light stain in the nervous system, is seen in the EHDP-exposed planarian. Anterior of the worms (brain) is to the top of the image. Scale bars: 0.5 mm.
